# Opposing immunomodulatory effects of the *Alternaria* mycotoxin tenuazonic acid in immune and intestinal epithelial cells

**DOI:** 10.64898/2026.06.24.734282

**Authors:** Vanessa Partsch, Francesco Crudo, Doris Marko

**Affiliations:** University of Vienna, Faculty of Chemistry, Department of Food Chemistry and Toxicology, 1090 Vienna, Austria; University of Vienna, Faculty of Chemistry, Doctoral School in Chemistry, 1090 Vienna, Austria

**Author notes:** **Corresponding author**: Dr. Francesco Crudo, Department of Food Chemistry and Toxicology, University of Vienna, Währinger Str. 38, 1090 Vienna, Austria.

**Keywords:** *Alternaria alternata*, food contaminants, immunomodulation, NF-κB signaling, intestinal epithelial cells, cytokines

## Abstract

Tenuazonic acid (TeA) is one of the most frequently detected *Alternaria* mycotoxins in contaminated food. Despite its frequent occurrence, its immunomodulatory effects remain insufficiently characterized. Therefore, the present study investigated the impact of TeA on inflammatory signaling and cytokine regulation in monocytes and intestinal epithelial cell (IEC) models. NF-κB activity was assessed using a reporter gene assay in THP1-Lucia™ monocytes, while cytokine mRNA expression and protein secretion were quantified in Caco-2 and HCEC-1CT cells by qRT-PCR and ELISA, respectively.

In THP-1 monocytes, TeA significantly suppressed lipopolysaccharide (LPS)-induced NF-κB activation in a concentration-dependent manner starting at 25 μM, while cytotoxicity occurred only at concentrations ≥100 μM. In HCEC-1CT and differentiated Caco-2 cells, TeA increased IL-6, IL-8, and TNF-α mRNA levels at non-cytotoxic concentrations (≥10 μM). In Caco-2 cells, these transcriptional changes were accompanied by increased cytokine secretion, whereas HCEC-1CT cells showed only partial effects on the protein level after short-term exposure. Following prolonged incubation, TNF-α secretion was increased and IL-6 and IL-8 secretion were slightly reduced. IL-10 remained unaffected under all conditions.

Overall, TeA exerted cell type-dependent immunomodulatory effects characterized by immunoinhibitory activity in monocytes and pro-inflammatory responses in IECs, highlighting the complex immunotoxic potential of this *Alternaria* mycotoxin.

## Introduction

Inflammatory signaling is a central component of immune defense, enabling rapid cellular responses to environmental stressors such as pathogens and dietary xenobiotics (Marshall et al. 2018). A key regulator of these processes is the NF-κB signaling pathway, which controls the transcription of major inflammatory mediators including interleukin (IL)-6, IL-8, and tumor necrosis factor-alpha (TNF-α). While tightly regulated NF-κB activation is essential for immune homeostasis, its dysregulation can contribute to inflammatory disorders and disease development (Liu et al. 2017).

Innate immune cells, including monocytes and macrophages, play a central role in initiating and regulating inflammatory responses to xenobiotics. In parallel, the intestinal epithelium represents the primary barrier for orally ingested food contaminants and actively contributes to immune regulation through cytokine secretion and signaling crosstalk. Disruption of these processes may affect both local and systemic immune homeostasis (Bouhet und Oswald 2005).

Diet represents a major route of exposure to bioactive secondary metabolites produced by fungi, including mycotoxins from *Alternaria* species (Schmutz et al. 2019). These ubiquitous plant pathogens contaminate a wide range of agricultural commodities during cultivation and storage, resulting in chronic dietary exposure to their toxins through commonly consumed foods. *Alternaria* mycotoxins comprise structurally diverse compounds, including dibenzo-α-pyrones such as alternariol (AOH), alternariol monomethyl ether (AME), and altenuene (ALT), perylene quinones such as alterperylenol (ALTP) and altertoxin I (ATX-I), and structurally distinct compounds including altersetin (AST) and tentoxin (TEN) (Aichinger et al. 2021). While many of these metabolites have been extensively investigated for their genotoxic and cytotoxic properties, increasing evidence suggests that they may also interfere with immune-related signaling pathways (Louro et al. 2024). In particular, several *Alternaria* mycotoxins have been reported to modulate NF-κB-dependent inflammatory responses. For example, AOH, AME, ALT, ALTP, ATX-I, AST, and TEN have been shown to suppress NF-κB activation in monocytic and macrophage cell models. In addition, AOH and ALTP have been reported to alter cytokine secretion profiles in immune and intestinal epithelial cells (Crudo et al. 2024; Kollarova et al. 2018; Partsch et al. 2025; Partsch et al. 2026; Schmutz et al. 2019)

Given the high abundance of tenuazonic acid (TeA) (Fig. 1), a tetramic acid derivate, in many naturally contaminated food commodities and agricultural products (EFSA, 2016), this study focused on elucidating its immunomodulatory potential. NF-κB activity was assessed in THP-1 Lucia™ monocytes using a reporter-based assay. In parallel, cytokine expression profiles (IL-6, IL-8, IL-10, TNF-α) were quantified in IL-1β-stimulated Caco-2 and HCEC-1CT intestinal epithelial cells, with selected findings validated at the protein level by ELISA.

**Fig. 1.**
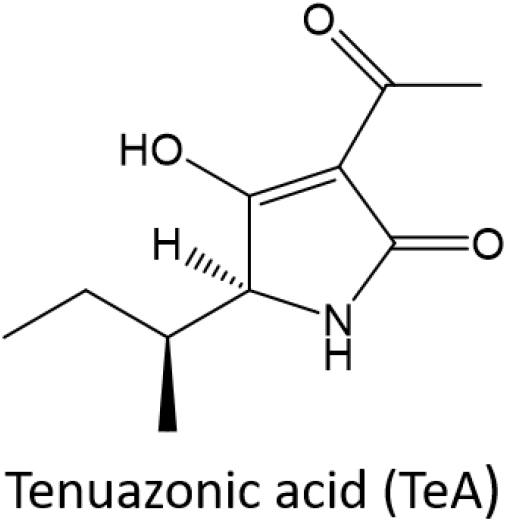
Chemical structure of the *Alternaria* mycotoxin tenuazonic acid (TeA)

## Materials and Methods

### Materials

Cell culture media and supplements were purchased from Thermo Fisher Scientific Inc. (Waltham, USA), Invivogen (San Diego, USA), Sigma Aldrich (St. Louis, USA), Merck KGaG (Darmstadt, Germany), Life Technologies Corporation (Carlsbad, USA) and Corning Inc. (Corning, USA). Lipopolysaccharide (LPS) from *Escherichia coli* and dexamethasone (Dexa) were acquired from Sigma Aldrich (St. Louis, USA), and the recombinant human IL-1β and the Quanti-Luc™ reagent from Invivogen (San Diego, USA). The CellTiter-Blue^®^ (CTB) reagent was purchased from Promega Corp. (Fitchburg, USA). The RNeasy Mini Kit and the primers for IL-6, IL-8, IL-10 and TNF-α were obtained from Qiagen N.V. (Hilden, Germany), while the primer for GAPDH from Eurofins Scientific SE (Luxembourg City, Luxembourg). The High-Capacity cDNA Reverse Transcription Kit, the *Power* SYBR™ Green PCR Master Mix, PCR plates and ELISA Kits (Human IL-6, IL8, IL-10 and TNF-α; uncoated) were purchased from Thermo Fisher Scientific Inc. (Waltham, USA). The *Alternaria* mycotoxin TeA (sc-202357A; LOT # G0225) was purchased from Santa Cruz Biotechnology, Inc. (Dallas, USA). Compound purity was declared to be >95% by the supplier and was verified by NMR analysis, which determined a purity of 98%.

### Cell culture

THP1-Lucia™ monocytes (InvivoGen, San Diego, USA) were cultured in RPMI 1640 medium supplemented with 10% heat-inactivated fetal bovine serum (FBS), 25 mM HEPES, 1% penicillin/streptomycin (P/S), and 100 μg/mL Normocin^®^. Reporter stability was maintained by culturing cells under selection pressure using 100 μg/mL Zeocin^®^ every second passage. Caco-2 brush border-expressing cells (C2BBe1, ATCC^®^ CRL-2102™) were cultured in DMEM (4.5 g/L glucose) supplemented with 10% FBS, 1 mM sodium pyruvate, 0.01 mg/mL insulin-transferrin-selenium (ITS), and 1% P/S. HCEC-1CT cells were kindly provided by Prof. Jerry W. Shay (UT Southwestern Medical Center, Dallas, USA) and maintained in DMEM supplemented with 2% cosmic calf serum, 2% Medium 199 (10×), 20 mM HEPES, 50 μg/mL gentamicin, 1 μg/mL hydrocortisone, 0.01 mg/mL ITS and 20 ng/mL human recombinant epidermal growth factor (EGF). All cell lines were maintained at 37 °C in a humidified atmosphere with 5% CO_2_ and passaged twice per week at ∼80% confluency.

### Cell seeding and treatment conditions

THP1-Lucia™ monocytes were seeded at a density of 3.5 × 10^5^ cells/cm^2^ in and used immediately after seeding. Caco-2 cells were seeded at 8.5 × 10^4^ cells/cm^2^ and differentiated for 7 days, with medium exchange every 2-3 days. HCEC-1CT cells were seeded at 1.7 × 10^4^ cells/cm^2^ and cultured for 48 h to form confluent monolayers.

Cells were exposed to different concentrations of TeA, ranging from 1-250 μM in THP-1 monocytes for the CTB and NF-κB assays. In IECs, concentrations of 1-100 μM for the CTB assay and 1-50 μM for qRT-PCR and ELISA were applied, respectively. To assess the immunosuppressive properties of the mycotoxin and mimic an inflamed environment, cells were pre-incubated with TeA for 2 h, after which THP-1 cells were stimulated with 10 ng/mL LPS and IECs with 25 ng/mL IL-1β. NF-κB and CTB assays were carried out with a total incubation time of 20 h (2 + 18 h). qRT-PCR and ELISA experiments were performed in IECs after 5 h (2 + 3 h) incubation, while in HCEC-1CT cells, an additional longer incubation period of 20 h was included.

### CellTiter-Blue^®^ (CTB) assay

The CTB assay was performed in all cell models to assess cytotoxicity and to define non-cytotoxic concentrations of TeA for subsequent qRT-PCR analyses in Caco-2 and HCEC-1CT cells. Cells were seeded in 96-well plates and incubated with TeA as described in the section “Cell seeding and treatment conditions”. As solvent control served DMSO (0.25% in THP-1 monocytes; 0.1% in IECs), and as the negative control 1 μM Dexa. After a total incubation period of 20 h, the CTB reagent was added at a 1:10 dilution and cells were incubated for an additional 2 h. In THP-1 cells, the reagent was diluted directly into the cell suspension, whereas in IECs, the incubation medium was removed and replaced with CTB reagent diluted in DMEM. Cells treated with Triton X-100 for 2 h (0.01% in THP-1 cells and 0.1% in IECs) served as positive controls for cytotoxicity. Fluorescence was measured at 560/590 nm (λex/λem) using a Cytation 3 microplate reader (BioTek Instruments, Winooski, USA) with the Gen5 software (v3.08).

### NF-κB reporter gene assay

To evaluate the effects of TeA on NF-κB signaling, the NF-κB reporter gene assay was performed in THP1-Lucia™ NF-κB monocytes as described by Partsch et al. (2026). Briefly. cells were treated under the same conditions as described for the CTB assay. To elucidate immunoinhibitory effects LPS was added after 2 h to active the pathway, while for the assessment of immunostimulatory effects cells were only incubated with TeA. After a total incubation time of 20 h, NF-κB activity was quantified according to the manufacturer’s instructions using Quanti-Luc™ reagent (coelenterazine-based substrate). Luminescence was measured using a microplate reader.

### Quantitative real time PCR (qRT-PCR)

Two-step qRT-PCR was performed to determine the effects of non-toxic concentrations of TeA on mRNA expression of IL-6, IL-8, IL-10, and TNF-α in Caco-2 and HCEC-1CT cells, as previously described by Partsch et al. (2026) Incubations were carried out in 24-well plates for Caco-2 and 12-well plates for HCEC-1CT cells. Experimental conditions were identical to those described for the CTB assay, with incubation times of 5 h for Caco-2 cells and 5 h and 20 h for HCEC-1CT cells. At the end of the incubation period, supernatants were collected and stored at -80 °C for subsequent ELISA analysis. Cells were washed with ice-cold PBS, and total RNA was isolated using the RNeasy Mini Kit (Qiagen, Hilden, Germany) according to the manufacturer’s instructions. RNA concentration and purity were assessed using a NanoDrop 2000 spectrophotometer (Thermo Fisher Scientific Inc., Waltham, USA).

For each sample, 1 μg RNA was reverse transcribed into cDNA using the High-Capacity cDNA Reverse Transcription Kit. qRT-PCR was performed with the Power SYBR™ Green Master Mix on a QuantStudio 3 system (Thermo Fisher Scientific Inc., Waltham, USA) in 96-well plates. Gene expression was analyzed for GAPDH (forward: TTCCCGTTCTCAGCCTTGAC, reverse: GATTTGGTCGTATTGGGCGC), IL-6, IL-8, IL-10, and TNF-α (Qiagen QuantiTect assays: Hs_IL6_1_SG, Hs_CXCL8_1_SG, Hs_IL10_1_SG, Hs_TNF_1_SG). Amplification conditions consisted of an initial activation step at 95 °C for 15 min, followed by 40 cycles of denaturation at 94 °C for 15 sec, annealing at 55 °C for 30 sec, and extension at 70 °C for 30 sec. Specificity was confirmed by melting curve analysis. Relative gene expression was calculated using the 2^−ΔΔCT^ method, with results normalized to GAPDH and expressed as fold change relative to the positive control (25 ng/ml IL-1β).

### Enzyme-linked immunosorbent assay (ELISA)

To evaluate whether TeA affects cytokine secretion in Caco-2 and HCEC-1CT cells, supernatants collected after treatment (see “Quantitative real time PCR”) were analyzed by ELISA. Protein levels of IL-6, IL-8, and TNF-α were quantified using commercial kits according to the manufacturer’s instructions.

### Statistical analysis

Statistical analysis was performed using OriginPro^®^ 2022 software (OriginLab, Northampton, USA). All experiments were conducted in at least three biological replicates, with technical triplicates for CTB and NF-κB reporter assays and technical duplicates for qRT-PCR and ELISA. Data were normalized to the respective positive control and are presented as mean + standard deviation (SD) of biological replicates. Statistical significance between TeA and the positive control was assessed using Student’s *t*-test. Differences among multiple concentrations of TeA were analyzed by one-way ANOVA followed by Fisher’s LSD post-hoc test.

## Results

### Immunosuppressive and cytotoxic effects on THP-1 monocytes

The immunosuppressive potential of the *Alternaria* mycotoxin TeA was investigated in THP-1 monocytes using the NF-κB reporter gene assay. Cells were exposed to seven concentrations of TeA (1–250 μM) in the presence or absence of 10 ng/mL LPS to assess immunoinhibitory and -stimulatory effects respectively. As shown in Fig. 2a, TeA significantly suppressed LPS-induced NF-κB activation in a concentration-dependent manner, starting at 25 μM (p < 0.001). To exclude cytotoxicity-related artefacts, cell viability was assessed in parallel using the CTB assay (Fig. 2b). A significant reduction in viable cells was observed at concentrations ≥ 100 μM (p < 0.001). Across the tested concentration range TeA did not induce immunostimulatory effects (see Supplementary Fig. 1).

**Fig. 2.**
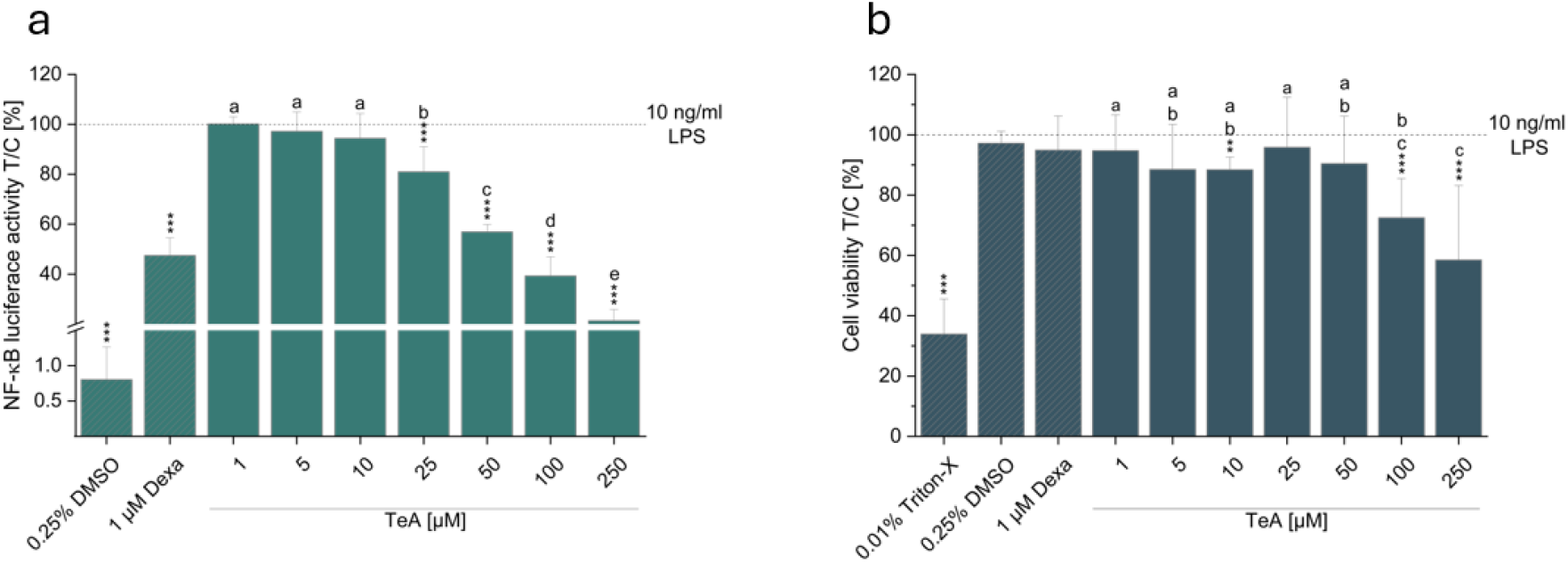
Immunoinhibitory and cytotoxic effects of the *Alternaria* mycotoxin tenuazonic acid (TeA) in THP1-Lucia™ monocytes. Panel a shows the results of the NF-κB reporter gene assay performed under co-stimulation with 10 ng/mL lipopolysaccharide (LPS). 1 μM dexamethasone (Dexa) served as a negative control and 0.25% DMSO as solvent control. Panel b shows cell viability assessed by the CellTiter-Blue^®^ assay. Triton X-100 (0.01%) was used as a positive control for cytotoxicity. Data are presented as mean + SD of at least three independent experiments and are expressed relative to the positive control (10 ng/mL LPS), as indicated by dotted lines. Statistical significance between treatments and the positive control was assessed using Student’s *t*-test (*p < 0.05, **p < 0.01, ***p < 0.001). One-way ANOVA followed by Fisher’s LSD post hoc test was applied to evaluate differences between different concentrations of TeA (a–e; p < 0.05)

### Impact of TeA on IL-6, IL-8, IL-10 and TNF-α gene transcription and secretion in Caco-2 and HCEC-1CT cells

To investigate the effects of TeA on IL-1β-induced cytokine gene transcription and secretion in HCEC-1CT and differentiated Caco-2 cells, mRNA and protein levels of IL-6, IL-8, IL-10, and TNF-α were quantified by qRT-PCR and ELISA, respectively. To ensure that only non-cytotoxic concentrations were analyzed, CTB assays were performed prior to gene expression analysis.

As shown in Fig 3, TeA induced cytotoxic effects in both cell lines starting at 50 μM (Caco-2, -p < 0.05; HCEC-1CT - p < 0.001). However, at 50 μM, cell viability remained at or above 80% in both IEC models.

**Fig. 3.**
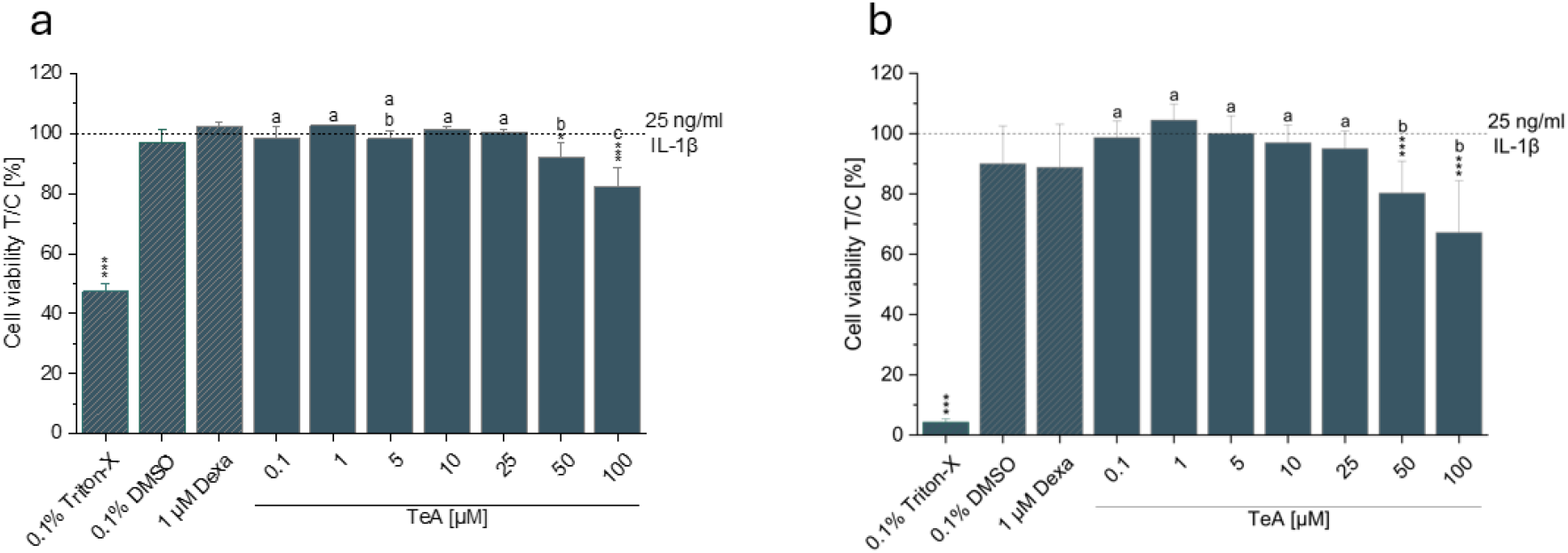
Cytotoxic effects of the *Alternaria* mycotoxin tenuazonic acid (TeA) on Caco-2 (a) and HCEC-1CT (b) cells. Cell viability was assessed by applying the CellTiter-Blue^®^ assay. Cells were pre-incubated with TeA or 1 μM dexamethasone (Dexa; used as a negative control) for 2 h, followed by co-stimulation with 25 ng/mL IL-1β for an additional 18 h. Triton X-100 (0.1 %) was used as a positive control for cytotoxicity. Data are presented as mean + SD of at least three independent experiments and are expressed relative to the positive control (25 ng/mL IL-1β), as indicated by dotted lines. Statistical significance between treatments and the positive control was assessed using Student’s *t*-test (*p < 0.05, **p < 0.01, ***p < 0.001). One-way ANOVA followed by Fisher’s LSD post hoc test was applied to evaluate differences between different concentrations of TeA (a–e; p < 0.05)

The assessment of cytokine regulation in Caco-2 cells after 5 h incubation revealed that TeA significantly increased IL-6 and IL-8 mRNA levels starting at 10 μM, whereas TNF-α expression was induced from 25 μM onwards (Fig. 4a). On the protein level, secretion of all three cytokines was significantly increased at 50 μM TeA (Fig. 4d).

**Fig. 4.**
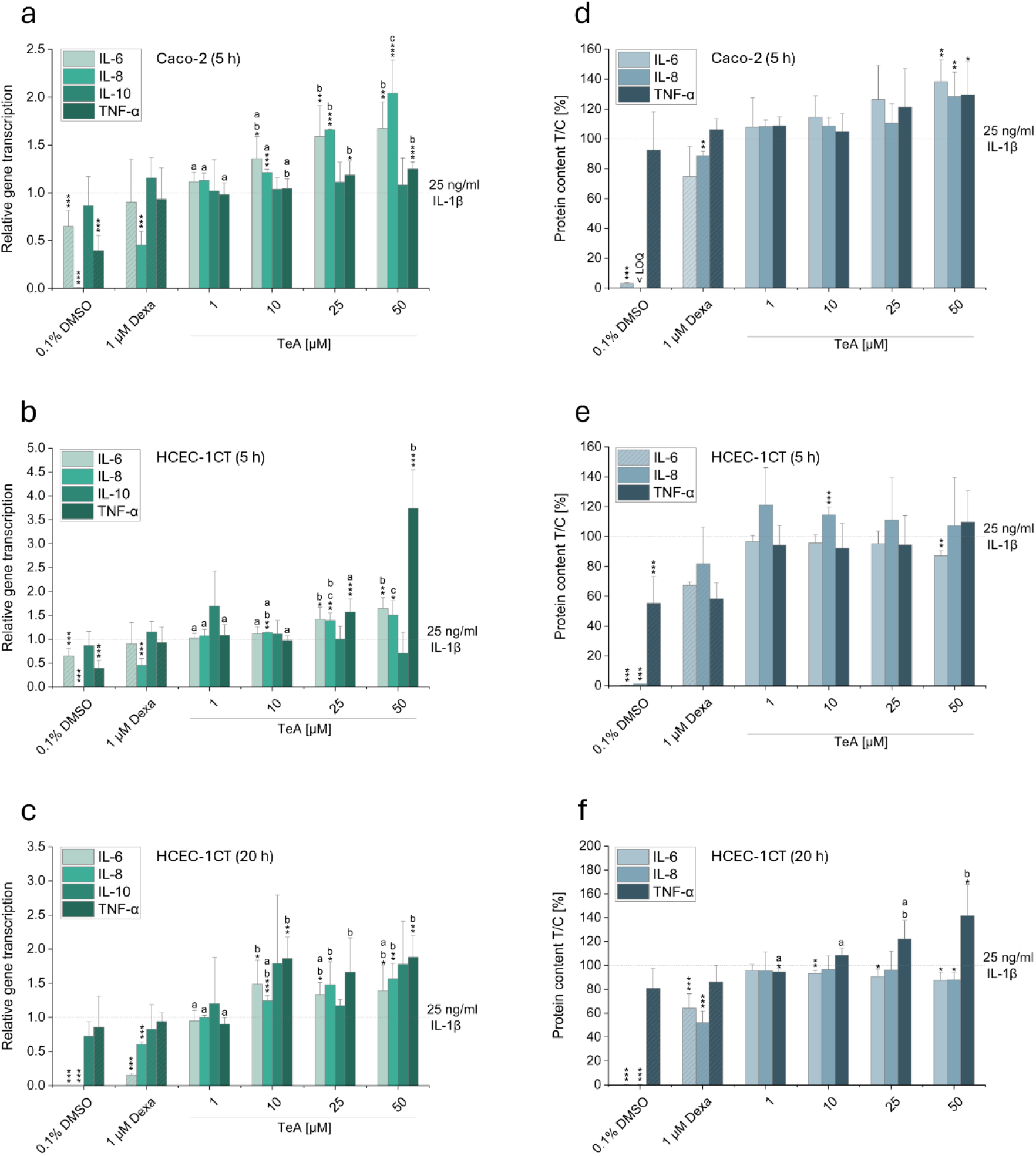
Impact of the *Alternaria* mycotoxin tenuazonic acid (TeA) on IL-6, IL-8, IL-10, and TNF-α gene transcription and protein secretion in IL-1β-stimulated Caco-2 (a,d) and HCEC-1CT (b,c,e,f) cells. Cells were pre-incubated with non-cytotoxic concentrations of TeA or 1 μM dexamethasone (Dexa; used as a negative control) for 2 h, followed by co-stimulation with 25 ng/mL IL-1β for an additional 3 h (a,b,d,e) or 18 h (c,f). Panels a–c show relative gene expression determined by qRT-PCR using the 2^−ΔΔCT^ method and normalized to GAPDH. Panels d–f depict corresponding cytokine secretion levels measured by ELISA. Results were normalized to the positive control (25 ng/mL IL-1β), as indicated by dotted lines. Data are presented as mean + SD of at least three independent experiments. Statistical significance between treatments and the positive control was assessed using Student’s *t*-test (*p < 0.05, **p < 0.01, ***p < 0.001). One-way ANOVA followed by Fisher’s LSD post hoc test was applied to evaluate differences between TeA concentrations (a–e; p < 0.05)

In HCEC-1CT cells, TeA significantly increased IL-8 mRNA levels at concentrations ≥10 μM and IL-6 and TNF-α levels at ≥25 μM after 5 h incubation. Notably, 50 μM TeA induced TNF-α transcription by more than 3.5-fold (Fig. 4b). These transcriptional changes were not reflected on the protein level, as only IL-6 secretion was slightly decreased at 50 μM (Fig. 4e). Since protein translation and secretion require longer incubation periods than transcriptional responses, HCEC-1CT cells were additionally exposed to TeA for 20 h. A transcriptional pattern comparable to that observed after 5 h was detected, with increased IL-6, IL-8, and TNF-α mRNA levels at concentrations ≥ 10 μM (Fig. 4c). Following prolonged incubation, TNF-α protein secretion was increased, whereas IL-6 and IL-8 secretion were slightly reduced (Fig. 4f). IL-10 transcription remained unaffected in both cell lines under all experimental conditions and was therefore not further analyzed on the protein level.

## Discussion

TeA is among the most frequently detected *Alternaria* mycotoxins in food (Aichinger et al. 2019). However, knowledge regarding its immunomodulatory properties remains limited (Louro et al. 2024). Therefore, the present study investigated its effects on NF-κB signaling and cytokine regulation in immune and intestinal epithelial cell models.

In THP-1 monocytes, TeA significantly inhibited LPS‐induced NF‐κB activation in a concentration‐dependent manner starting at 25 μM, with cytotoxic effects only observed at concentrations ≥100 μM (Fig. 2). These findings suggest that TeA can exert immunosuppressive effects under the experimental conditions applied. A previous study did not observe effects on NF‐κB signaling at the concentrations that were active in the present study (Crudo et al. 2024). Variations between studies may relate to material‐specific factors, as commercially available mycotoxins are often isolated from fungal cultures and may therefore differ in purity. While no additional purity assessment beyond the supplier’s declaration was performed in the previous study, preventing a direct evaluation of potential material‐related differences, the identity and purity of the TeA batch used in the present study were confirmed by NMR analysis, which indicated a purity of 98%. Nevertheless, the present results are consistent with previous reports showing that several structurally distinct *Alternaria* mycotoxins can suppress inflammatory responses and interfere with NF-κB signaling in THP-1 monocytes and macrophages (Crudo et al. 2024; Kollarova et al. 2018; Partsch et al. 2025; Partsch et al. 2026). Collectively, these observations suggest that different *Alternaria* mycotoxins can modulate innate immune signaling pathways despite their structural diversity. Given the central role of NF-κB in inflammatory responses, such inhibition may impair the ability of immune cells to appropriately respond to immunological and microbial stimuli (Liu et al. 2017).

Immune responses to xenobiotics are not solely regulated by immune cells but are also influenced by regulatory signals originating from IECs. As the first cellular barrier to orally ingested compounds, IECs play a dual role in both restricting toxin passage and actively communicating with underlying immune cells. This bidirectional crosstalk is essential for maintaining mucosal homeostasis, and its disruption has been associated with dysbiosis, chronic inflammation and tumor development (Yao et al. 2024). To investigate whether TeA affects the immune responsiveness of IECs, cytokine expression and secretion were examined in IL-1β-stimulated Caco-2 and HCEC-1CT cell models. In contrast to the immunoinhibitory effects observed in monocytes, TeA induced predominantly pro-inflammatory responses in the intestinal cell models. In differentiated Caco-2 cells, TeA increased transcription of IL-6 and IL-8 starting at 10 μM and TNF-α from 25 μM onwards (Fig. 4a). This was accompanied by elevated secretion of all three cytokines at 50 μM (Fig. 4d). Similar transcriptional effects were observed in HCEC-1CT cells (Fig. 4b,c). However, short-term exposure (5 h) resulted only in limited changes on the protein level, suggesting delayed translation or post-transcriptional regulation (Fig. 4e). Following prolonged incubation, increased TNF-α secretion was observed, whereas IL-6 and IL-8 secretion were slightly reduced (Fig. 4f). These findings indicate a discrepancy between transcriptional and protein-level responses to TeA, which may further depend on incubation time and cell type.

Furthermore, the contrasting responses observed between immune cells and IECs highlight the cell-type-specific nature of TeA-mediated immunomodulation. While inhibition of NF-κB signaling in THP-1 monocytes reflects an immunosuppressive effect, the increased cytokine expression detected in IECs points toward a pro-inflammatory response at the intestinal barrier. Comparable cell type-dependent effects were also observed for AOH, which was reported to induce immunosuppressive responses in monocytes and macrophages but upregulated TNF-α transcription in Caco-2 and HCEC-1CT cells, accompanied by increased TNF-α secretion in Caco-2 cells (Partsch et al. 2026; Schmutz et al. 2019). These divergent responses likely reflect the distinct physiological roles and basal inflammatory states of the respective cell types. Monocytes are primed for rapid and robust inflammatory signaling, whereas IECs are characterized by tightly regulated immune responses that are essential for maintaining intestinal barrier integrity and preventing excessive immune activation (Marshall et al. 2018; Wullaert et al. 2011).

Overall, the present results demonstrate that TeA modulates inflammatory signaling in a cell-type-dependent manner. Furthermore, the study expands current knowledge regarding the immunotoxic potential of *Alternaria* mycotoxins. These findings highlight the importance of further mechanistic studies to identify the molecular targets of TeA.

## Data availability

The datasets generated during and/or analyzed during the current study are available from the corresponding author on reasonable request.

## Author contributions

Conceptualization: Doris Marko, Francesco Crudo; Methodology: Francesco Crudo, Vanessa Partsch; Formal analysis and investigation: Vanessa Partsch; Writing - original draft preparation: Vanessa Partsch; Writing - review and editing: Doris Marko, Francesco Crudo; Funding acquisition: Doris Marko; Supervision: Doris Marko, Francesco Crudo

## Abbreviations

ALT: Altenuene
AOH: Alternariol
AME: Alternariol monomethyl ether
ALTP: Alterperylenol
AST: Altersetin
ATX-I: Altertoxin I
CTB: CellTiter Blue
Dexa: Dexamethasone
ELISA: Enzyme-linked immunosorbent
IL: Interleukin
ICE: Intestinal epithelial cell
LPS: Lipopolysaccharide
NF-κB: Nuclear factor kappa-light-chain-enhancer of activated B cells
qRT-PCR: Quantitative real time PCR
TEN: Tentoxin
TeA: Tenuazonic acid
TNF-α: Tumor necrosis factor-alpha

