## Supplementary Figure 1 for "Opposing immunomodulatory effects of the *Alternaria* mycotoxin tenuazonic acid in immune and intestinal epithelial cells"

**Corresponding author:**

Dr. Francesco Crudo

ORCID:

Doris Marko: 0000-0001-6568-2944

Francesco Crudo: 0000-0002-4876-8057

Vanessa Partsch: 0009-0000-4073-7013

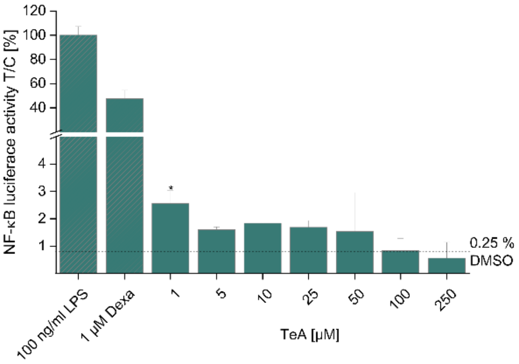


**Supplementary Fig. 1** Immunostimulatory effects of the *Alternaria* mycotoxin tenuazonic acid (TeA) assessed by applying the NF-κB reporter gene assay.

THP1-Lucia™ monocytes were exposed to TeA for 20 h. Cells incubated with 1 µM dexamethasone (Dexa) for 2 h followed by co-stimulation with 10 ng/mL lipopolysaccharide (LPS)for 18 h served as a negative control and just LPS as positive control. Data are presented as mean + SD of at least three independent experiments and are expressed relative to the solvent control (0.25% DMSO), as indicated by a dotted line. Statistical significance between treatments and the solvent control was assessed using Student's *t*-test (*p < 0.05, **p < 0.01, ***p < 0.001).
